# Centriole and transition zone structures in photoreceptor cilia revealed by cryo-electron tomography

**DOI:** 10.1101/2023.10.05.560879

**Authors:** Zhixian Zhang, Abigail Moye, Feng He, Muyuan Chen, Melina A. Agosto, Theodore G. Wensel

## Abstract

Primary cilia mediate sensory signaling in multiple organisms and cell types but have structures adapted for specific roles. Structural defects in them lead to devastating diseases known as ciliopathies in humans. Key to their functions are structures at their base: the basal body, the transition zone, the “Y-shaped links” and the “ciliary necklace”. We have used cryo-electron tomography with subtomogram averaging and conventional TEM to elucidate the structures associated with the basal region of the “connecting cilia” of rod outer segments in mouse retina. The longitudinal variations in microtubule (MT) structures and the lumenal scaffold complexes connecting them have been determined, as well as membrane-associated transition zone structures: Y-shaped links connecting MT to the membrane, and ciliary beads connected to them that protrude from the cell surface and form a necklace-like structure. These results represent a clearer structural scaffold onto which molecules, identified genetics, proteomics, and superresolution fluorescence, can be placed in our emerging model of photoreceptor sensory cilia.

**Summary:** Cryo-electron tomography and subtomogram averaging reveal new structural features at the base of the light sensing cilia of retinal rods. These include the basal body, the Y-links between axoneme and membrane, and the ciliary necklace of the transition zone.

## Introduction

Photoreceptors are sensory neurons, which sense and transduce light signals into changes in electrical current and neurotransmitter release. To perform this task robustly, these cells have developed highly-ordered compartmentalization and high rates of inter-compartmental protein trafficking. Proteins required for phototransduction and the structural integrity of the light-sensing outer segment (OS) compartment and the stacks of membranous discs it contains are transported from the inner segment (IS), where biosynthesis primarily occurs, through a thin and crowded intermediate compartment known as the connecting cilium (CC). In each rod cell of the mouse retina, roughly 60 rhodopsin molecules per second are synthesized and packaged into OS membrane discs (1, 2), along with hundreds of other proteins transported to the OS. About 80 new discs are formed per day in each rod. Consequently, this trafficking structure and system must be tightly regulated and well-maintained.

The photoreceptor CC and OS together form a unique, specialized sensory organelle with modified central core templated from the ubiquitous eukaryotic signaling hub, the primary cilium (reviewed in (3)). Primary cilia are cylindrical organelles extending from the cell surface that are involved in extracellular sensing and signaling. The cilia are packed with a distinctive set of lipids and proteins, including receptors, ion channels, microtubule-associated structural proteins, and proteins associated with a complex and tightly regulated bi-directional trafficking machinery (Berbari et al., 2009; Hilgendorf et al., 2016; Klena and Pigino, 2022; Nachury and Mick, 2019; Pazour and Witman, 2003). Cilia generally form after cells exit the cell cycle, as the primary, immotile cilia do in rod and cone photoreceptors after their last cell division. Some features of the photoreceptor sensory cilium and its earliest stages of development are similar to those in other ciliated cells, whereas others are unique to photoreceptors (Baehr et al., 2019; May-Simera et al., 2017; Wensel et al., 2021). Ciliogenesis initiates when the centrioles migrate to the apical edge of the inner segment (IS) and form a basal body, consisting of a mother centriole, from which the axoneme eventually grows, and an adjacent daughter centriole, both embedded in a matrix of amorphous appearance but complex organization (Fry et al., 2017; Mennella et al., 2014; Sonnen et al., 2012) known as pericentriolar material (PCM). The centrioles contain 9 symmetrically arranged microtubule triplets (MTTs) at one end (referred to here as the proximal end), which extend and transition into 9 MT doublets (MTDs) at the distal end, from which the axoneme grows (Kumar and Reiter, 2021; Li et al., 2019). These MTDs continue throughout the ∼1.2 μm CC, transitioning into singlets as it extends up the ∼20 μm long OS. At the distal end of the CC (the base of the OS), the light-sensing discs are formed. The region at the base of cilia where the microtubules undergo a transition from triplets to doublets is often referred to as the transition zone (TZ), and its unique complement of proteins is thought to regulate trafficking through the region, performing a “gate” function (Park and Leroux, 2022). In mouse rods, only a subset of TZ proteins are restricted to the base of the CC, with most of those found beyond the triplet-to-doublet transition, whereas others, such as CEP290 (Potter et al., 2021) and SPATA7 (Dharmat et al., 2018) are distributed throughout the entire length of the CC, suggesting the entire CC may contribute to the gate function. In addition, some ciliary proteins such as centrins (Trojan et al., 2008) and EB1 (Hidalgo-de-Quintana et al., 2015) are found in the CC, rather than restricted to the basal body, as in most other primary cilia.

There are multiple inherited human diseases, known as ciliopathies, due to mutations in genes encoding cilium-or basal-body associated proteins and resulting in defects in cilium structure and function. In many cases, these multi-syndromic diseases include retinal degeneration (Bachmann-Gagescu and Neuhauss, 2019; Braun and Hildebrandt, 2017; Chen et al., 2021; Seo and Datta, 2017), and in a number of cases, retinal degeneration is the only major symptom (Estrada-Cuzcano et al., 2012), highlighting the importance of the BB and CC in photoreceptor function. To better understand how these disorders can cause photoreceptor degeneration, deep knowledge of the structures of these compartments is indispensable.

In this study, we used cryo-electron tomography to determine structures at nanometer-scale resolution of the BB, CC, and associated structures in rod photoreceptors of mice. These findings, and their integration with the growing body of structural, genetic, and biochemical data on cilia and BB from many other tissues and species (Dang et al., 2017; Danielsson et al., 2020; Dean et al., 2016; Li et al., 2004; Liu et al., 2007; Pazour, 2004; Sun et al., 2019), could help provide insight into the roles of these molecular machines and their components in photoreceptor function and health. Hopefully these findings will add to the understanding needed for therapies designed to protect or restore visual function in retinal ciliopathies.

## Results

### Structural domains identified from mouse photoreceptor centrioles and connecting cilium

To facilitate studies of the basal body centrioles (hereafter referred to as centrioles) and connecting cilium, fragments of rods, including rod outer segments (ROS) with attached distal rod inner segments (RIS) were isolated and imaged by cryo-electron tomography (cryo-ET) as described previously (Gilliam et al., 2012; Robichaux et al., 2019; Wensel and Gilliam, 2015). The method employed to isolate ROS allows for retention of the entire connecting cilium (CC) and basal body (mother and daughter centriole). Fig. 1A displays a partially segmented 50 nm slice from a reconstructed tomogram, with the mother (M) and daughter (D) centrioles and CC highlighted in different colors to demonstrate the direction and position of each centriole and CC. The cylinder-shaped basal body has an average length of 400 nm. The proximal region (0-170 nm, light magenta) contains MTTs, the mid-region (170-340nm, blue) has incomplete triplets transitioning to doublets, and the distal region (340-400nm, light violet) contains MTDs. Note that the transition to doublets occurs well within the centrioles and not in the CC that emerges beyond the ciliary pocket, as was documented previously in mammalian centrioles (Le Guennec et al., 2020; Sun et al., 2019). The doublets then continue longitudinally into the CC, as shown in cyan in Fig. 1A. Cross-sectional slices of the tomograms are shown in Fig. 1B and the corresponding subtomogram average density maps (see details of the averaging below) of the MTTs, incomplete triplets, centriolar MTDs, or CC MTDs are shown in Fig. 1C, using the same color scheme. Sub-tomogram averaging (Fig. 1C; see discussion below) allows clear visualization of individual tubulin subunits (13 A-tubule protofilaments, 10 B- and C-tubule protofilaments, consistent with what is seen in other basal bodies and centrioles (Guichard, 2013; Li, 2012; Agard, 2019)), as well as non-tubulin proteins both on the inside (microtubule inner proteins, MIPs, Figs. 1C, 1D, discussed below) and outside (Pinhead, A-C linker, and other external densities) of the MTs.

**Figure 1.**
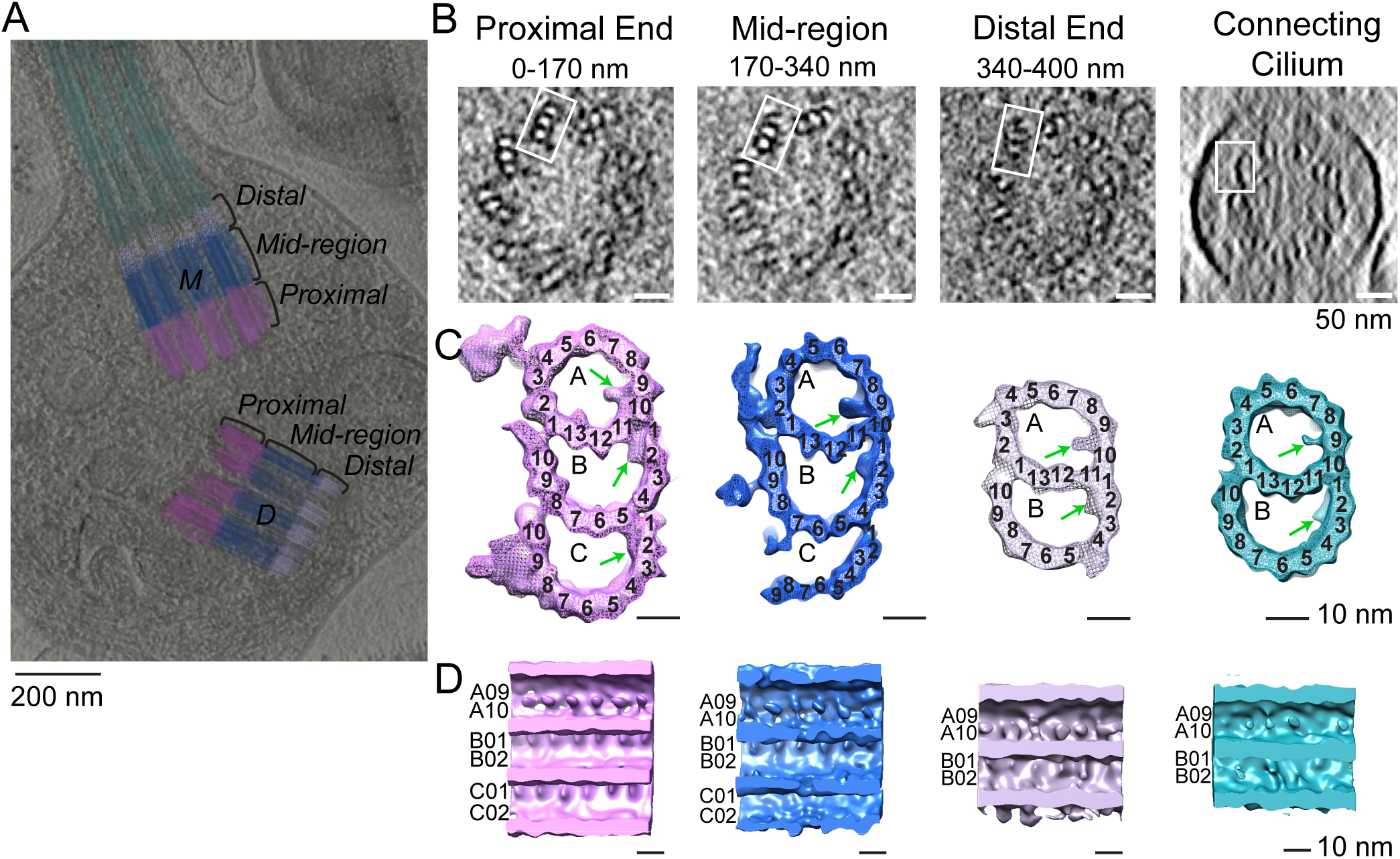
Tomographic reconstruction and subtomogram averaging of three structural domains from mouse centrioles. (A) A slice from a reconstructed tomogram showing mother and daughter centrioles (denoted ‘M’ and ‘D’). Centrioles were partitioned longitudinally into three regions, the proximal end (0-170 nm, light magenta) consisting of MT triplets, the mid-region (170-340 nm, blue) consisting of incomplete triplets, and the distal end (340-400 nm, light violet), consisting of MT doublets. The extending CC doublets are denoted in cyan. (B) Slices of a tomogram of a daughter centriole through the three centriole regions and the CC (from a different part of the tomogram). White boxes indicate examples of approximate regions used for initial stages of subtomogram averaging and refinement. (C) Four maps were generated by averaging tomogram sub-volumes in each region of centrioles and of the CC. The A-, B-, and C-tubules, including non-tubulin densities, are shown, with protofilament numbering. Microtubule inner proteins (MIPs) are shown with green arrows. (D) Maps in C cut in half and rotated 90° toward viewer to show microtubule inner proteins (MIPs).

### Models of Centriole and Connecting Cilium

In many cases, the cilia and basal body tomogram reconstructions appear elliptical in cross-section, rather than perfectly circular, due to flattening and compression during grid preparation. To improve the signal-to-noise ratio and resolution of the centriolar structure, unflattening and averaging by imposition of 9-fold symmetry, as described previously (Robichaux et al., 2019), was applied to selected centriole maps that displayed minimal flattening. Regions of the tomograms containing doublet, triplet or incomplete triplet MTs were boxed out of raw tomograms in both mother and daughter centrioles and used for subtomogram averaging (Chen et al., 2019) over multiple individual BB and CC. No consistent differences were observed between the mother and daughter centriole maps except for the extended region of doublets forming the CC axoneme extending from the distal end of the mother centriole, so both were used for subtomogram averaging. Separate sub-maps were used for CC doublets and distal BB doublets. The averaged maps were fit into the symmetrized map of either the centriole or the CC. Fig. 2A displays the 9-fold symmetry-imposed maps for the proximal, mid, and distal regions of the BB. Superimposed on the right side of each of these maps are subdomain structures of the MTT, incomplete MTT, or MTD obtained by subtomogram averaging. These were fitted into the symmetry-averaged maps, as shown in the models in Fig. 2B, revealing the individual tubulin subunits and non-tubulin structures associated with the centriole. This model, defined in more detail in Methods, was used for further analysis.

**Figure 2.**
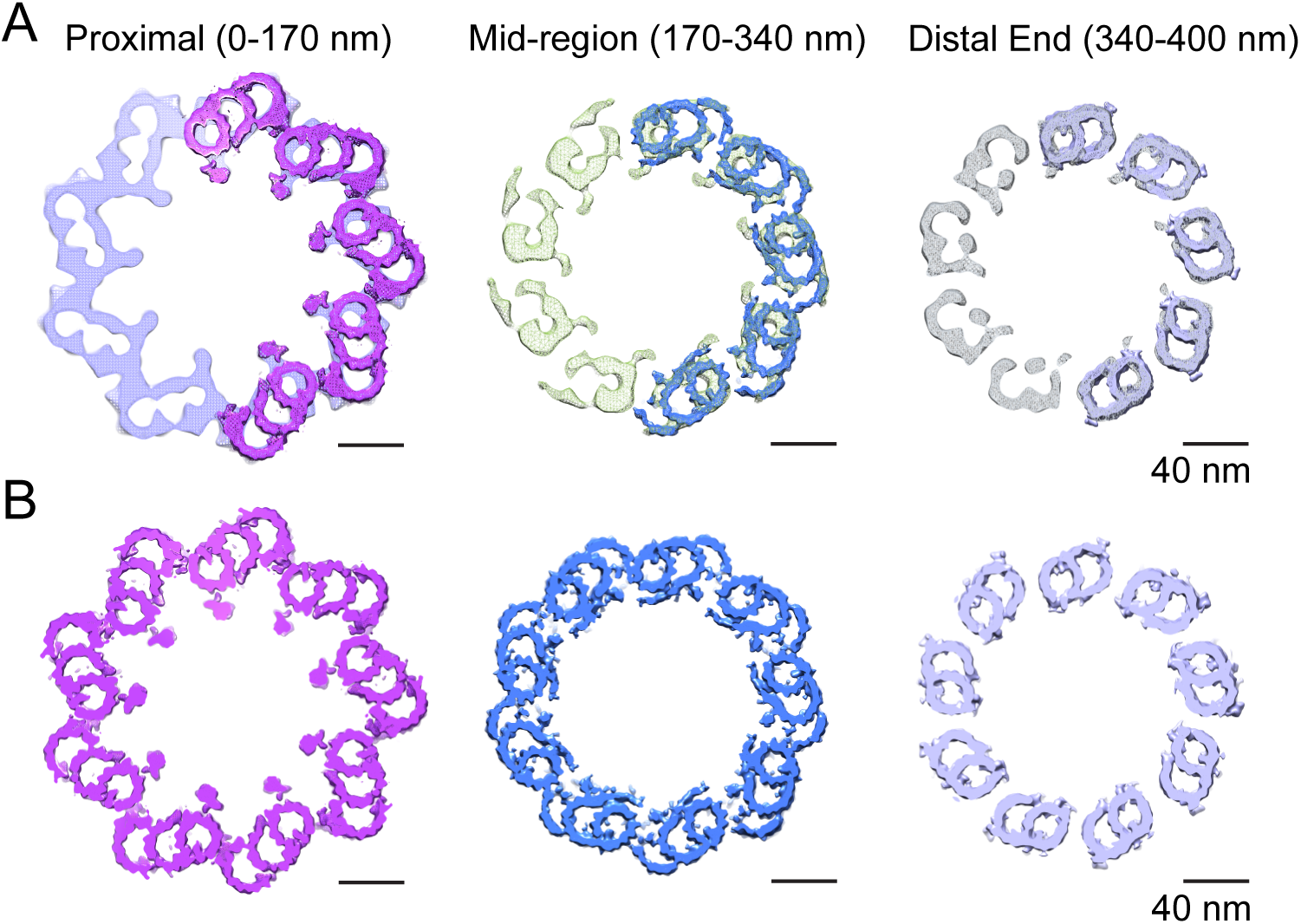
Centriole model. (A) 9-fold symmetrized cross-sections of a map generated by rotational averaging of a daughter centriole at the proximal (*left*), middle (*middle*) and distal (*right*) regions, respectively, with maps generated by subtomogram averaging of triplet, incomplete triple or doublet averages (those shown in Fig 1C) fitted onto the symmetrized map and superimposed on the right side of each corresponding map. (B) The fully refined, symmetrized reconstructed/fitted models used for examination of extra-MT densities and inter-MT connections for MTTs and MTDs.

### Twist Angles Observed Throughout Basal Body and Connecting Cilium

There are two distinct types of “twist” associated with the arrangement of MTs along the long axis of the centriole and CC. To aid in visualizing these, we fit the average MT maps into raw maps of a centriole (Fig. 3) or CC (Supp. Fig. S1). One type of twist (Anderson, 1972; Sun et al., 2019) involves the angle of the long axes of the MTDs and MTTs as they wrap helically around the long central axis of the MT bundle. As shown in Supplemental Fig. S1, the outermost (C for triplets, B for doublets) protofilament forms an imperfect right-handed helix with a very long repeat period (>400 nm), such that one period is not complete along the centriole. The lack of multiple repeats, the variation in this twist angle in the CC *vs.* that in the centriole, the switch from *C* to *B* MTs as the outermost MT during the transition from triplet to doublets, and bending of the CC (Fig. 3A) make it difficult to assign a precise number to this period.

**Figure 3.**
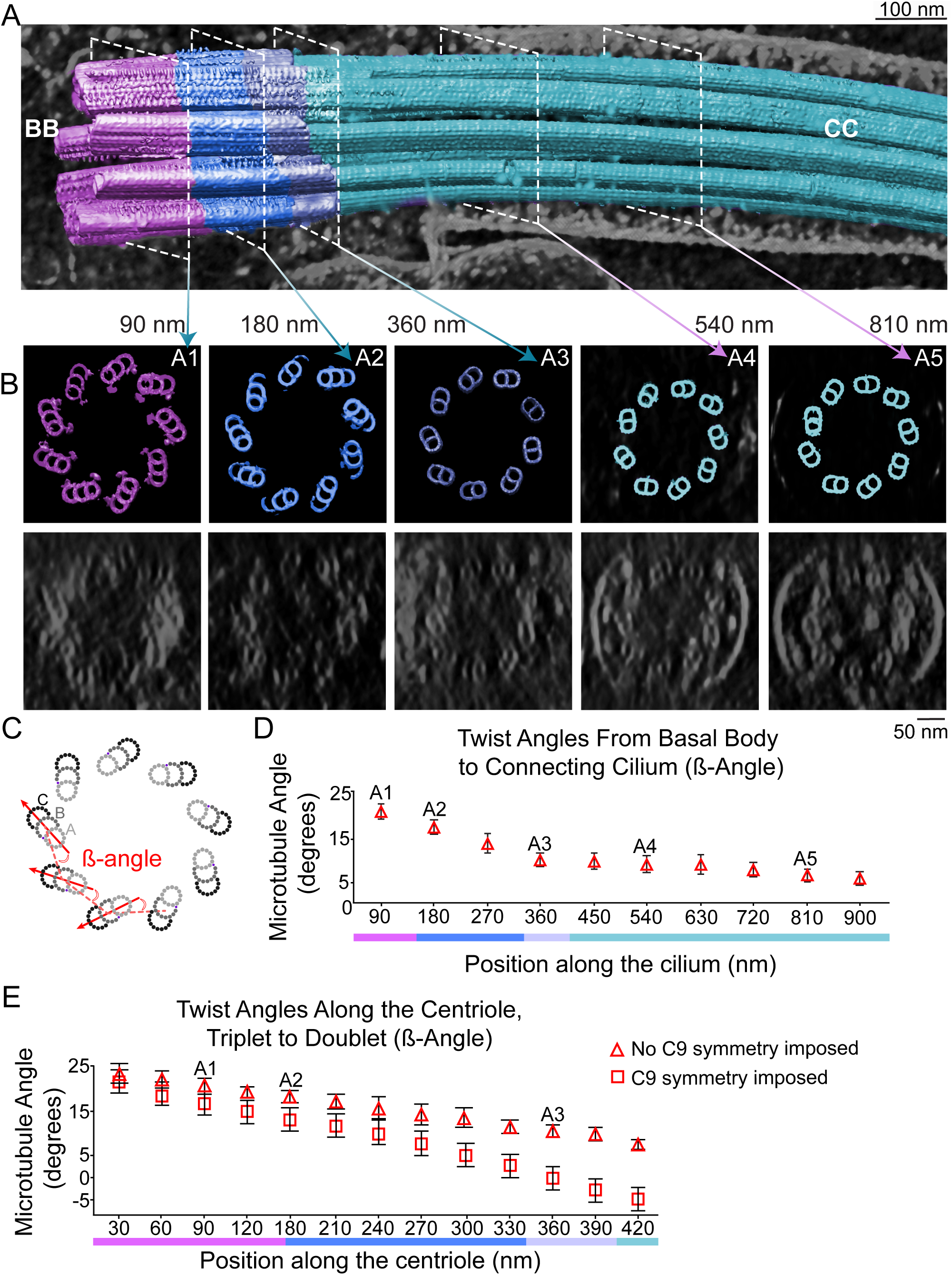
Fit of MT subtomogram averages into raw tomogram and range of MT twist angles in the cross-sectional plane. (A) The averaged sub-volume of triplets (cyan; the same sub-volumes used for Figure 2) and doublets (magenta) were fitted back into the raw tomogram of an entire mother centriole-CC to generate a whole-cilium map. (B) Cross sections (top panel, model, bottom panel raw tomogram) corresponding the positions indicated by dashed white lines in A and denoted by A1-A5 in panel B. The twist angles β (defined in panel C) were measured and averaged for each cross-section and plotted in D for the cilium map and E for the centriole map. as a function of longitudinal position. This procedure was repeated using a 9-fold rotationally averaged centriole map (Supplementary Fig. S1) and plotted as boxes in E. Points and error bars represent means ± S.D.

A similar gradual twist can be observed in recent reconstructions of bovine motile cilia (Greenan et al., 2020) and primary cilia of epithelial cells (Sun et al., 2019), and was also reported in an early study of monkey oviduct cilia (Anderson, 1972), albeit as a left-handed twist; note that ambiguity in projection direction during reconstruction can lead to flipping of handedness (Anderson, 1972; Chen et al., 2019). A cryo-ET study of primary cilia from MDCK cells revealed a consistent right-handed twist of MT in the basal portion of the cilium of 56.4° ± 7.3° per μm, that became highly variable in the distal portions (Kiesel et al., 2020). The width of the MT bundle also steadily declines along the proximal-to-distal axis in our maps, ranging in diameter from 230 nm at the proximal (complete triplet) end to 190 nm at the distal end (doublets), due to both the second type of twist described below and to the loss of the C MT.

The second type of “twist” previously reported in the literature (Anderson, 1972; Le Guennec et al., 2020; Li et al., 2012; Paintrand et al., 1992), and most readily visualized in cross-sectional views, refers to the angle the row of each MTT or MTD forms with respect to the radius of the 9 + 0 bundle, or equivalently (but with a different value), the angle each triplet or doublet makes with respect to the line connecting it to the adjacent MT row in the bundle (Fig. 3C, Supplemental Fig. S1D). We measured this “twist” using two different angles (Fig. 3D and Fig. S1D). One, the *α* angle (Greenan et al., 2020), is defined as the outer angle between the lines connecting the centers of A- and B-tubules within each doublet or triplet and the lines connecting the A-tubules of adjacent triplets or doublets, as shown in Fig. S1D. The *β* angle *(Li et al., 2012)* is defined as the inner angle between the lines connecting the centers of the B-tubules of each doublet or triplet and the line connecting the centers of the A- and B-tubules of the adjacent triplet or doublet (Fig. 3C). For 9-fold symmetric MT bundles, these two angles have a simple geometric relationship (see legend to Supplemental Fig. S1), with a nearly constant difference of ∼30° for the angles in our maps.

Both angles decrease from proximal to distal centriole ends (Fig. 3, Fig. S1D). The α-angle displays a steady decrease from a 51° MTT angular rotation in the proximal end down to a 25° MTD angular rotation in the CC (Fig. S1D). A similar range was observed in mammalian motile cilia (bovine tracheal epithelium, (Greenan et al., 2020)). The β-angle decreased from a 21° MTT angle in the proximal end down to a 6° MTD angular rotation in the distal end of the CC (Fig. 3D). Different numbers are obtained depending on whether the raw tomograms or symmetrized maps are used, due to distortions in the angles caused by both physical flattening and computational un-flattening (Fig. 3E). Measurement of the β-angle in raw maps of the centriole, without symmetrization, resulted in a 21° angle for the proximal MTTs, ∼18° for the incomplete MTTs in the mid-region, and an 11° angular rotation of the MTDs in the distal centrioles (Fig. 3E and Supplemental Fig. S1D). These angles differ somewhat when calculated from symmetrized maps (17° proximal, 13° mid, 0° distal Fig. 3E). Likely the correct values of these angles in intact retina lie somewhere in between those for symmetrized and non-symmetrized maps, but a definitive answer to this question will probably depend on obtaining data from samples obtained by high-pressure freezing and focused ion beam milling of more intact retina samples (Poge et al., 2021; Rigort et al., 2010; Young and Villa, 2023; Zhao et al., 2021).

### Non-tubulin complexes associated with centriole microtubules

In addition to the tubulin subunits and protofilaments of the MT, our maps and model have clearly visible features that are formed by proteins other than tubulin isoforms. These include microtubule inner proteins (MIPs) found within the lumenal regions of the MT and complexes on the outside of the MT such as the pinhead, and the inner scaffold complexes.

### Microtutubule Inner Proteins (MIPs)

Protrusions into the centriolar MT lumen, examples of a general class of microtubule inner proteins (MIPs), are seen in the A-(Figs. 1, 2, 4), B- and C-tubules (Fig. 4B), with the one in the A-tubule being most prominent. These are located at protofilaments (PF) A09/A10, B01/B02, and (with the weakest density) C01/02. MIPs have previously been reported in the centriole (Greenan et al., 2018), as well as determination of some of their molecular components (Fabritius et al., 2021; Gui et al., 2021; Ichikawa et al., 2019; Ichikawa et al., 2017; Kiesel et al., 2020; Li et al., 2022; Ma et al., 2019).

**Figure 4.**
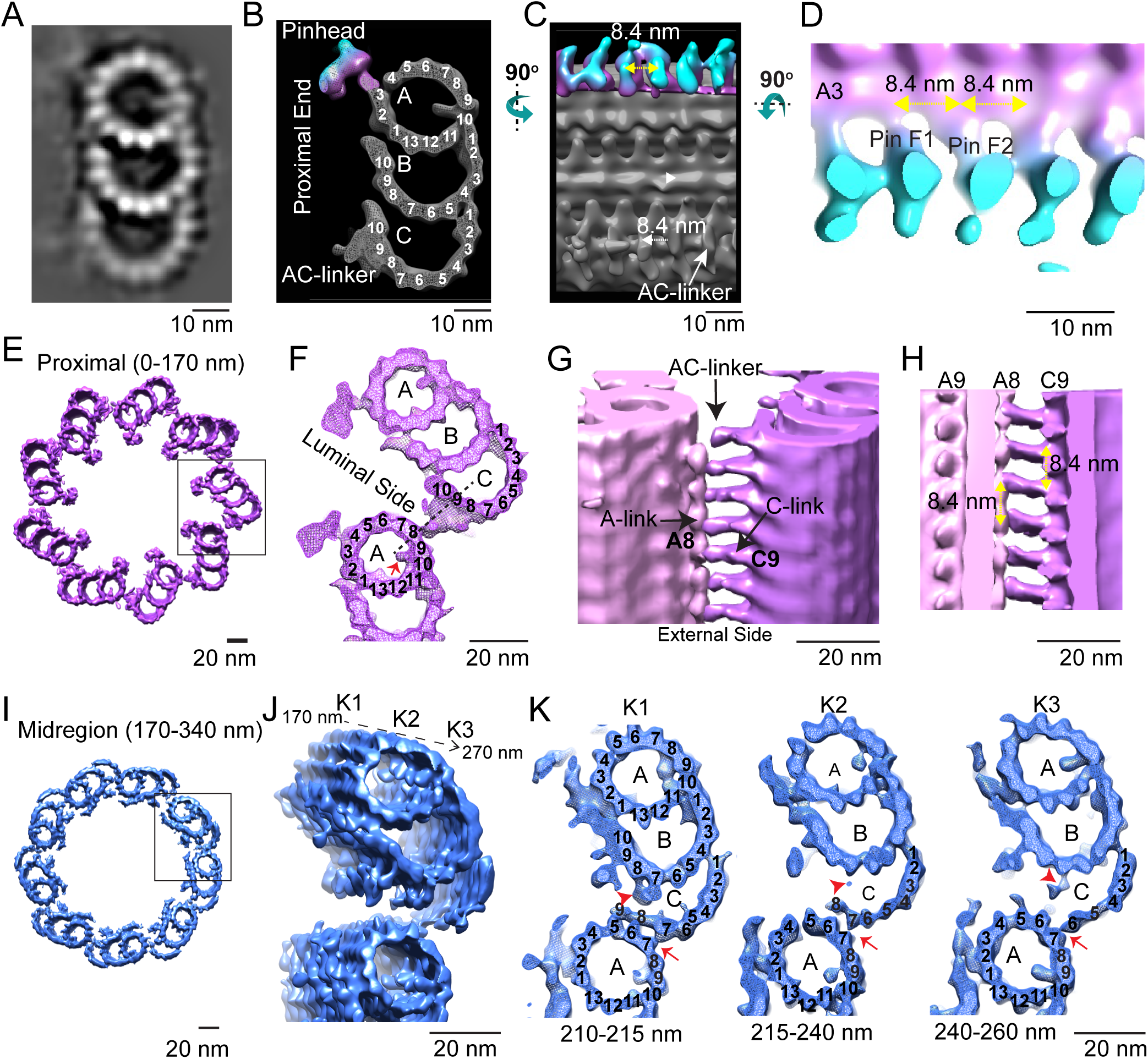
Pinhead and A-C linker in the proximal and mid-regions of the centriole. (A) Projection view and (B) surface view (with surface capping) of section through MTT map obtained from subtomogram averaging of complete MTTs in the proximal region (0-170 nm) viewed down the long axis of the MTT. (C) Surface rendering from B rotated 90° to reveal the side facing the lumen of the MT bundle. The pinheads are colored with depth-coding (magenta closer to MT, cyan furthest from the MT surface). (D) is a magnified and rotated version of the rendering in C, highlighting the Pin F1 and Pin F2 feet of the pinhead and their 8.4 nm spacing. (E) Surface rendering of centriole model in the proximal region of the mother centriole (0-170 nm) (from Figure 2B). (F) Magnified view of the boxed area in (E). The A-C linker is the density that links PF A8 on the A MT at the bottom to PF C9 of the adjacent C MT of the MTT at the top. (G) Side view from the outside of the MT bundle highlighting the A-C linkers. (H) Cut-away view (the cutting plane is shown as a dotted line in F) from the lumen of the MT bundle with yellow arrows highlighting the 8.4 nm spacing of the A-C linkers. (I) Mid-region (170 nm to 340 nm) of the compete centriole model showing the incomplete MTTs; the proximal 100 nm of the boxed region is shown at higher magnification in J and K. (J) is a tilted version showing the progression of structural changes along the centriole axis. (K) shows cross-sectional views of three different sub-regions (K1-K3, extracted at different levels from the map as shown in J. Red arrows and arrowheads indicate the variations in the connections between A MT and C MT along the axis.

### Pinhead

One of the conserved structures within the proximal region of the basal body is the pinhead, which protrudes from PF A3 (protofilament 3 of the A-tubule; see numbering in Fig. 4B) into the basal body lumen (Fig. 4A, B; Fig.Supplemental Figure S2). The unattached end of the pinhead splits into two densities, or “feet” (PinF1 and PinF2). The longitudinal spacing between each pinhead’s PinF2 and the subsequent pinhead’s PF1, which is the same as that between PinF1 and PinF2 of a single pinhead, is 8.4 nm (Fig. 4C, D). Although this centriolar structure is conserved throughout species, there have been differences reported in the spacing between pinhead feet; *e.g.*, 8.2 nm in CHO cell centrioles (Greenan et al., 2018), and 8 nm in *Chlamydomonas* (Li et al., 2019). There has also been a report in some *Trychonympha* species of a difference in spacing between feet in each pinhead (7.9 nm) as compared to spacing between adjacent pinheads (8.6 nm) (Nazarov et al., 2020) (Fig. S2). The molecular constituents of the pinhead are not known; CEP135 has been proposed as a major component, but a definitive determination has not been made (reviewed in (Gonczy and Hatzopoulos, 2019)).

### Inter-microtubule connections in the basal body

Connections between adjacent MTT can be seen in our model and are most readily visualized in cross-sectional views (Fig. 4E). In the proximal (0-170 nm) region of complete triplets, the AC linker, a structure connecting the A- and C-tubules in adjacent triplets in this region can be seen (cross-sections, Fig. 4F-4H; longitudinal views from the outside, 4G, or inside, 4H of the MT bundle). The longitudinal spacing of the AC linkers is 8.4 nm, as for the pinhead. At the junction between PF C10 and PF B7, there is extra density associated with C10 (Fig. 4), which may be a non-tubulin subunit. There is a gradual change of these inter-MT connections in the mid-region (170-340 nm, Fig. 4I, tilted view, Fig. 4J) of incomplete triplets. The gradual loss of the C MTs, first of PF C9 and C10, and eventually the remaining C PFs is apparent. There is a persistent, albeit weak density associated with PF B7, near the original position of C10, which may represent the non-tubulin protein at partial occupancy (Fig. 4K, red arrowhead). The gradual loss of the C MT is accompanied by a replacement of the AC linker connection with what appear to be direct connections of C9 to A6 and of C7 to A7. In the progression from triplets through incomplete triplets to doublets, there is first (210-215nm) a break at the inner junction of the C-tubule. The A-C linker is remodeled, and, in its place, the major attachments observed are initially between PF A5, A6 and A7 of the A-tubule, and C9, C8 and C7, respectively, (Fig. 4K, panel K1). As the C-tubule becomes further disassembled, the connections to the neighboring A-tubule are lost, with the C6-to-A7 connection being the last to remain (Fig. 4K, panels K2, 215-240 nm and K3, 240-260 nm).

The transitions of connections between the A-tubule and C-tubule along the distance from the proximal to the distal ends have also been observed in centrioles or basal bodies from other species and cell types with either motile or sensory cilia. In the CHO cell centriole, at the proximal A-C linker, PF A9 is connected to C8/C9; at the distal end, and PF A9 is directly bound to PF C8 (Greenan et al., 2018). In *Chlamydomonas reinhardtii*, at the proximal end, PF A6 is connected to the neighboring triplet at the tail end of the C-tubule; and at distal end of the BB, PF A6 is connected with PF C7/C8 (Li et al., 2012). It is unclear whether the proteins that make up the A-C linker change throughout this transition region, or if it is the same proteins that have remodeled to allow detachment and eventual loss of the C-tubule.

### Inner scaffold structure in basal body

Along the lumenal side of the MT bundle the presence of filamentous structures attached to the A- and B-tubule, termed Arm A and Arm B (Fig. 5), were observed throughout the mid-region (170-340 nm) and the distal region (340-400 nm) of the centriole (Fig. 5A, arrows). Together, these form the structure termed the inner scaffold. At the boundary between the proximal triplets and mid-region incomplete triplets, there is density in the Arm A region whose position is similar to some of the density observed in the pinhead. It is unclear if distinct molecular components give rise to these densities. Line intensity plots from the raw tomograms demonstrated that these filamentous densities exhibited 8.5 ± 1.1 nm spacing along the length of the microtubules (Fig. 5B, Fig. S3). Cross-sections of rotationally averaged maps also clearly show this inner scaffold in the mid- and distal centriole regions, as well as in the CC, compared to the proximal centriolar region (Fig. S4). Whether or not the proteins that form this inner scaffold change from centriole to CC is unclear. To enhance the structural detail for the inner scaffold, subtomogram averaging of the structure was carried out, followed by back-fitting into the centriole map from Fig. 2B. This reconstruction allowed for clear visualization of the two distinct densities: Arm A (attached to A2) and Arm B (attached to B9) (Fig. 5C, D, E F, indicated in orange). Both have a periodicity of 8.4-8.6 nm (Figs. 5B, 5F, 5G), similar to spacings in other cell types (Fig. S5). In cross-sections from the 3D reconstructions, these densities formed an almost continuous ring (Fig. 5H, lower panel), even though only partial inner scaffold filaments were observed in the raw tomogram due to sample compression and the missing wedge in tilt-series images of individual centrioles (Fig. 5H, upper panel). An additional density between Arm A and Arm B seen in some other centrioles (a “stem” in, *e.g.*, *P. tetraurelia*, (Le Guennec et al., 2020)) was not apparent in our maps (Fig. S6). In centrioles from other species and cell types (Le Guennec et al., 2020), this inner scaffold has been consistently observed, first appearing in the incomplete triplet region and extending over various distal lengths (Fig. S6).

**Figure 5.**
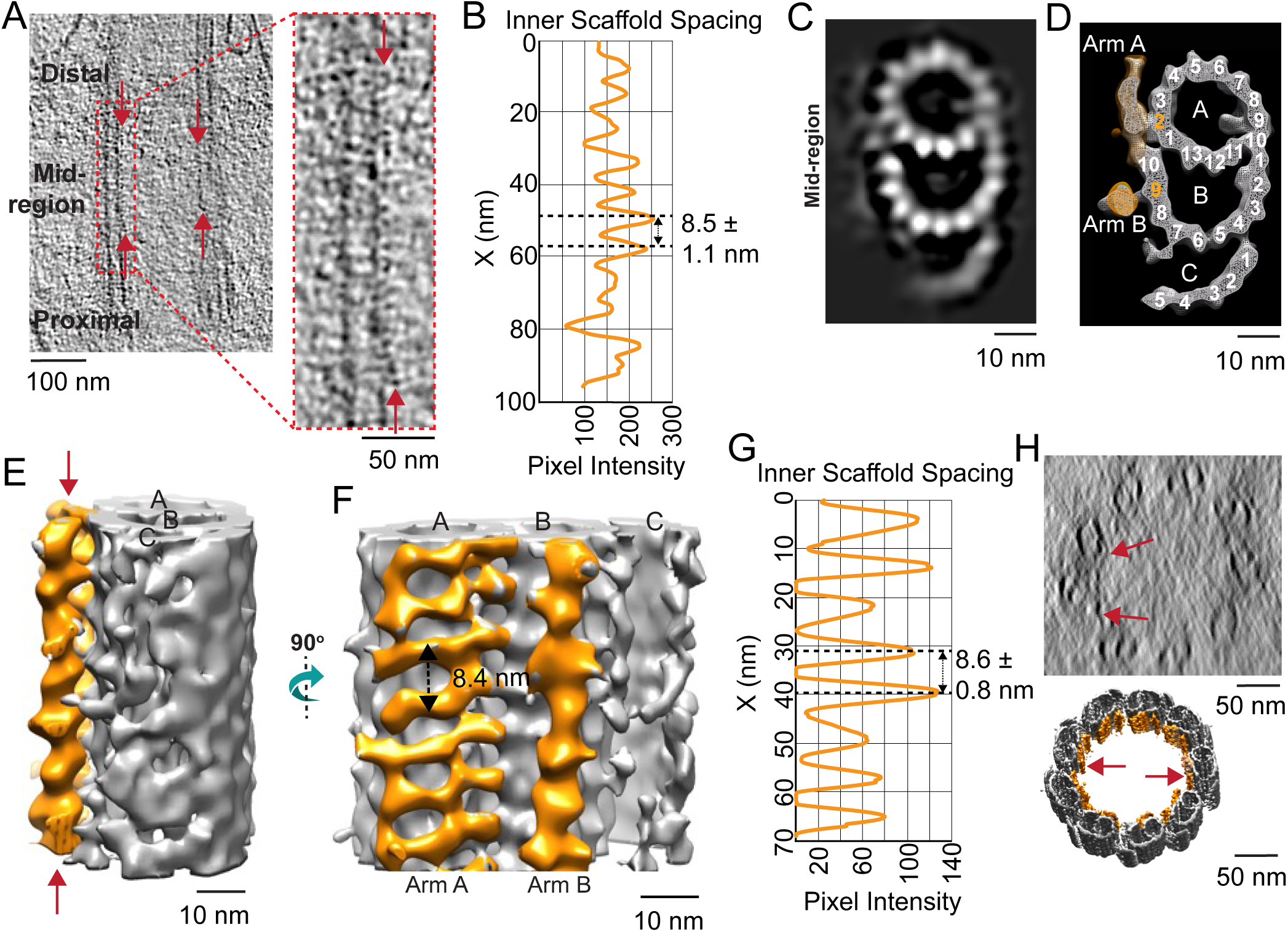
Inner scaffold along the incomplete triplets in the mid-region of the basal body. (A) Section from raw cryo-electron tomogram of rod photoreceptor showing the basal body structure in longitudinal view with labels for the proximal, mid-region and distal positions along the BB. The red arrows around the mid-region indicate the visible inner scaffold densities attached to the MTs. A zoomed-in view of inner scaffold densities attached to MTTs (as indicated with dashed red box) is shown to the right. (B) The corresponding line intensity profile (right) of the inner scaffold showing an average periodicity in the mid region of 8.5 nm. (C, D) Cross sections of the BB mid-region map from subtomogram averaging in projection view (B) or surface view with surface capping (D). These reveal the complete A-, B-, and partial C-tubules. In panel D, the extending filamentous densities from the lumen side of PF A2 (Arm-A) and PF B9 (Arm-B) are highlighted in orange. (E, F) Longitudinal views of the same region as in C and D, with inner scaffold shown in *orange* bracketed by red arrows in E. The view in F is of the map shown in E rotated 90° toward the viewer. The line intensity profile along the direction indicated by the double-headed arrow in F is shown in panel G; (periodicity of 8.6 nm). (H) Cross sectional view of a section of a raw tomogram in the BB mid-region with red arrows indicating presence of inner scaffold (top) and a slightly tilted view of the corresponding region of the model from Figure 2B highlighting the inner scaffold indicated in *orange* (bottom).

### Ciliary necklace beads and bridges to MT observed along the connecting cilium

Two of the hallmarks of the transition zone region at the base of cilia are 1) the presence of arrays of external protuberances from the ciliary membrane, known as the “ciliary necklace” (Gilula and Satir, 1972) and 2) connections between the MT and ciliary membrane known as “Y-shaped links” or “Y-links” due to their greater width at the membrane as compared to the narrow neck connected to the MT. Photoreceptor CC were among the first cilia identified as having a ciliary necklace, through freeze-fracture studies that implicated their association with intramembrane particles (Rohlich, 1975), observed throughout most of the length of the CC. Photoreceptor CC also have Y-links not only at their base, but throughout their length (Potter et al., 2021); however, neither the relationship between the Y-links and necklace beads nor their three-dimensional structure has been clearly determined, and their molecular compositions are also unknown.

Examples of the appearance of the ciliary necklace can be seen in raw tomograms (Fig. 6A), as well as in conventional TEM images (Fig. 6C). Ciliary necklace beads encircle the cilium as strands or rows along the circumference of the ciliary membrane, lying roughly in planes tilted slightly from those perpendicular to the ciliary axis, as seen in the surface rendering of the 3D map (Fig. 6D). The intramembranous strands are roughly parallel to one another, and many, if not all, appear to span only a portion of the circumference of the CC membrane.

**Figure 6.**
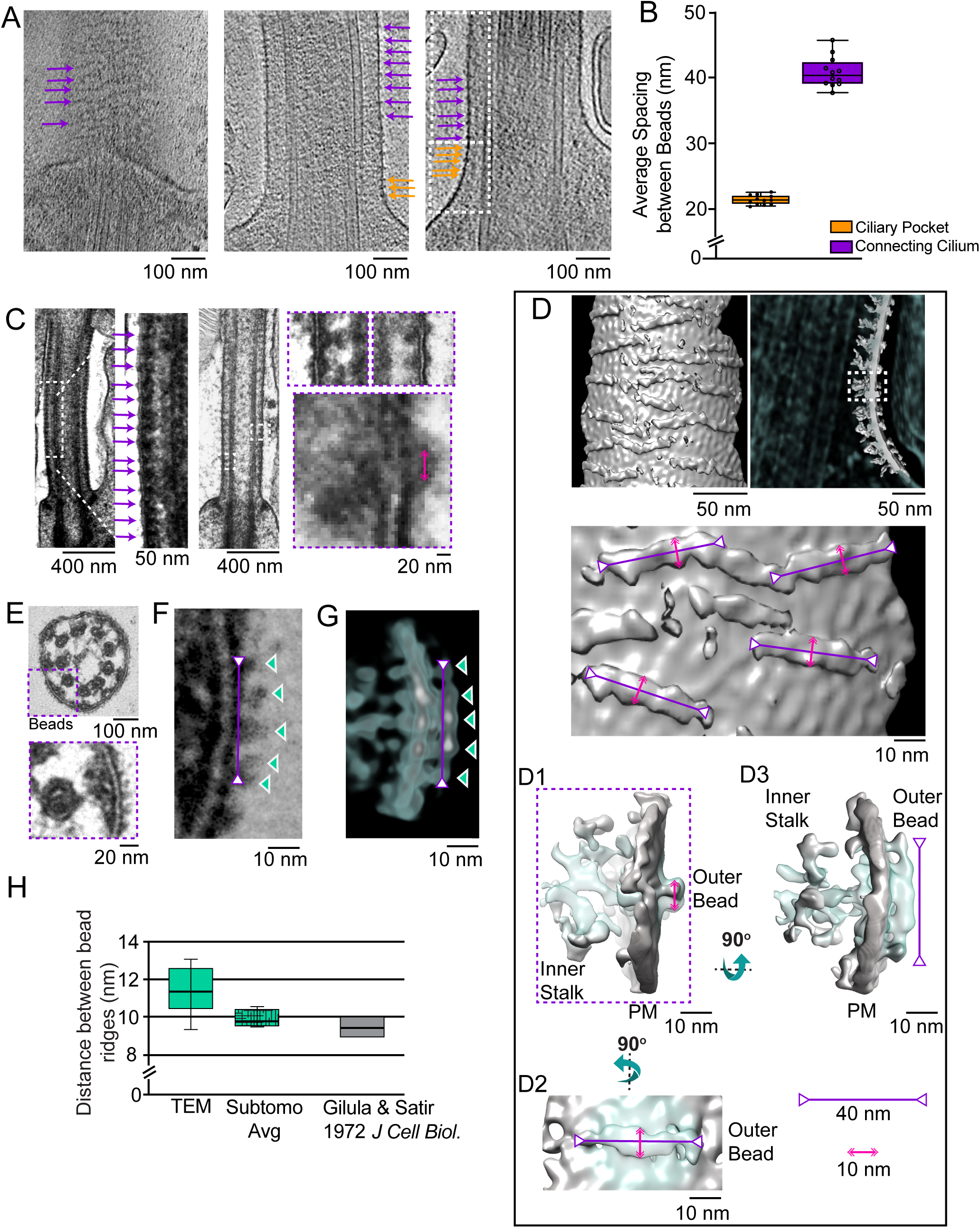
Subtomogram averaging of the intramembranous beads/ciliary necklace along ciliary membrane. (A) Sections from raw tomograms from mouse rod photoreceptors displaying intramembranous beads periodically arranged on the ciliary plasma membrane (left displaying a slice from the outside view of the CC; right showing a slice through the center of the CC). The beads were categorized into two groups based on spacing, ciliary pocket region (orange arrows, ∼22 nm spacing), and CC region (purple arrows, ∼40 nm). (B) Spacing between beads for each of the two groups along the ciliary membrane. (∼ 22 nm, n=6 for ciliary pocket and ∼40 nm, n =7 for CC. Box and whisker plot shows median (horizontal line), upper and lower quartile (box edges), and range of measurements (error bars), as well as individual points. (C) Conventional electron micrographs of mouse rod CC displaying the same intramembranous beads (average spacing 38.5 nm apart in this image, magenta arrows) along the CC. The dashed-line box in the image on the far left indicates the region show at higher magnification in the (middle) image immediately to its right. Dashed-line boxes on the image to the far right are the regions shown at higher magnification below. (D) The map of the ciliary necklace beads obtained by subtomogram averaging was fit (in multiple copies) into a raw tomogram of a CC and the resulting map shown in surface view from outside the CC (top, far left), in a projection view sliced through the CC membrane (immediately to the right of the surface view), and a magnified surface view (lower left). The series of images on the right show different high-magnification views of the average bead (cyan) and the portion of the CC membrane into which it was fit (gray). The longest dimension of the averaged structure is about 40 nm (magenta bars with inverted arrowheads) and the narrower dimension is about 10 nm (pink double-headed arrows). (E, F) Conventional TEM images of CC showing the beads. The dashed-line box in the upper image of E shows the region shown at higher magnifications below and in panel F, to the right. The magenta bar with two inverted arrowheads represents 40 nm, and the cyan-outlined arrowheads indicate the ridge-like subdomains of the structure that yield the “bead” appearance. G is a similar view of a portion of a map from cryo-ET obtained as described for panel D above. (H) Plot of spacings measured for bead ridges in our TEM images and subtomogram averages, or in a previously published report of freeze-fracture/SEM studies. Whisker plots show medians, lower and upper quartiles and range of the data.

Measurements of the longitudinal distances between adjacent beads along the length of the CC (Fig. 6B) revealed that the beads were ∼40 nm apart along most of the CC, whereas closer to the basal body (ciliary neck) the beads were ∼22 nm apart (n=7, Fig. 6B). The 40nm distance corresponds well to the spacing of intra-membranous particles seen in a previous freeze-fracture study of rat rods (Rohlich, 1975). Our measurements using conventional TEM images of ultrathin sections of mouse CC yielded an average bead spacing along the CC of ∼35nm (not shown), consistent with previous measurements from such images of ∼32 nm in rat rods (Besharse et al., 1985). The differences could reflect distortions arising from the extensive processing of the conventional TEM and freeze-fracture/SEM samples, as compared to the unfixed nature of the flash-frozen cryo-ET samples.

Volumes containing beads of the ciliary necklace were extracted and subjected to subtomogram averaging, followed by fitting of the average structure into the raw tomogram of a CC base (see Methods). In the resulting map (Fig. 6E), the ciliary necklace beads appear to contain two major domains connected within the ciliary membrane. An outer bead structure, ∼40 nm in the circumferential dimension (purple inverted-head arrows, Fig. 6D) and <10 nm in the longitudinal dimension (pink arrows, Fig. 6D), gives rise to the “necklace” appearance. An inner bridge is composed of multiple small subdomains, suggestive of a noisy average of heterogeneous structures filling the distance between the MT and the membrane (Fig. 6D right panel). The circumferential span of the outer bead structure appears to have 5 globular ridges, which are clearly distinguished in conventional TEM images (Fig. 6E, F) and in an averaged subtomogram map of such images (Fig 6G) but are barely resolved in the raw tomograms (Fig. 6A). The distance between these 5 ridges differs slightly between subtomogram average maps (∼11 nm, Fig. 6G, H) and conventional TEM images (∼10 nm; Fig. 6F, H), and is close to the measurements described from freeze fracture studies of rat rods (Rohlich, 1975).

### Y-link structures along the connecting cilium

When maps of MTDs with connected partial inner bridges/links and of beads are fitted into a 9-fold rotationally averaged CC map (Supplemental Fig. S7A and Fig. 7A, prepared by computational straightening, unflattening and rotational averaging of a single CC map, as described previously (Robichaux et al., 2019)), it is apparent that there is good alignment and partial overlap of the two structures (Fig. 7B). Within the CC, a narrow neck-like structure is connected at a somewhat variable position between PF A10 and PF B1, which extends and widens in the radial direction toward the ciliary membrane (Fig. 7A, B). The structure of this Y-link appears to be somewhat heterogenous, as is apparent in the variations between different longitudinal regions in the rotationally averaged maps (Fig. 7E), as well as in conventional TEM images (Fig. 7F). Because of this heterogeneity, the inner bridge/link portion of the “bead” map obtained by subtomogram averaging likely represents an average of multiple distinct structures rather than an accurate representation of a single structure. Despite the intrinsic differences in sample preparation referred to above, the cryo-ET and conventional TEM results are remarkably well-aligned (Fig. 7D). From both TEM and cryo-ET results, the widest part of the inner stalk (Y-link) was measured to be ∼41 nm. Integration of the results from raw tomograms, subtomogram averages, conventional TEM and prior literature leads to the model illustrated in Fig. 7G. In this model, the wide portion of the Y-links occupies a ∼15° arc of the CC circumference, so there is a 25° separation between adjacent Y-links, and there is continuous density leading from the MT doublet to and through the ciliary membrane, forming 5 globular ridges per Y-link.

**Figure 7.**
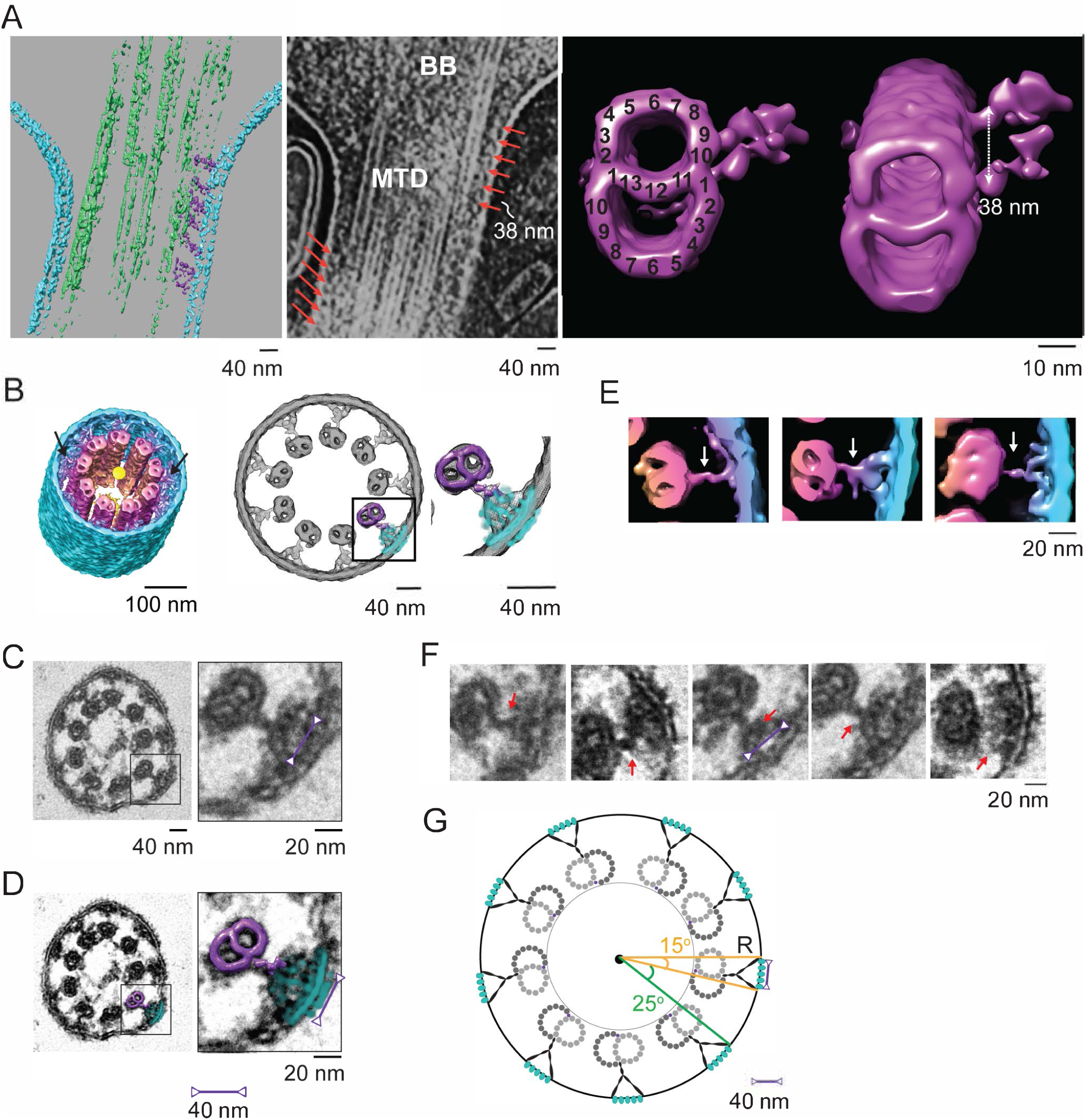
Y-link connections to microtubules. (A) *Left panel*, segmented version and *middle panel*, solid representation of tomogram used to generate the template for the subtomogram average of doublets with attached bridges/links (*right panel, magenta*). BB, basal body, MTD, microtubule doublets. The MT-to-membrane bridges/links are indicated with orange arrows. The subtomogram average map on the right displays longitudinal spacings of 38 nm (*double-headed arrow*), also seen in the raw tomogram (*orange arrows*). The attached bridging density emerging from the A/B junction (A10/B1) is consistent with the observation in the CC longitudinal view (*left* and *middle panels)*. (B) Rotationally averaged map of the CC shown in tilted (left) and cross-sectional (center) views; arrows point to Y-links. The averaged doublet + filamentous density map (magenta) and the averaged ciliary bead map (from Fig. 6, cyan) were fit into the rotationally averaged map of the CC (gray mesh), providing a composite model of the MTD-Y-link-membrane complex (square box in *center* panel and expanded view on the *right*). For comparison, in C, a conventional TEM image of a cross-sectional view of a detergent-extracted CC is shown at two different magnifications, with a magenta double-headed arrow displaying a distance of 41 nm. The MTD-Y-link ciliary model, aligned as in panel B is shown superimposed on the TEM image in D. (E) A view of the same map as in B, with the cross-section taken at a different position along the axis, and with distance from the center color-coded. Three individual MTD-Y-link-membrane-bead complexes at different axial positions are shown to the right, with white arrows indicating the bridging/link density. (F) Different cross-sectional images from conventional TEM of CC, with red arrows pointing to the filamentous densities. (G) schematic model of the MTD-Y-link-membrane-bead complexes in CC cross-section. The “Y” portion is meant to denote approximate position, rather than three-dimensional structure, given the heterogeneity demonstrated in E and F.

## Discussion

### Conserved and unique structural features of photoreceptor ciliary centrioles

The overall architecture of the centrioles associated with the photoreceptor sensory cilium is remarkably similar to those associated with other cilia, including both motile and primary cilia, and including those of protists metazoan organisms. However, many of the structural details we have observed can vary substantially among cell types and developmental stages and have not previously been determined for mammalian photoreceptors. Some of these details suggest a re-framing of standard models of these organelles.

### Longitudinal Microtubule twisting

The variation in MTT angle in the cross-sectional plane along the centriole axis is well-documented in multiple ciliated cells. The helical twist of the MT is perhaps less well appreciated, albeit also well documented. Reviews of ciliary or centriole structures tend to feature cartoons showing perfectly straight MT in both structures, absolutely parallel to the long axes. The functional significance of this twist is unclear, but it has been proposed (Sun et al., 2019) to help sensory cilia withstand mechanical stress due to flow. It is hard to imagine a need for this function in photoreceptors, as the close-packed nature of outer segments holds the cilia fairly rigidly in place. However, as the results of our method of preparing cell fragments for microscopy demonstrate, the photoreceptor CC and basal body are remarkably resistant to such forces. In those experiments the outer segments are broken off from the retina by shear forces, but the CC and BB consistently remain intact.

### Width and cross-sectional twist angle shifts in BB in other cell types

Although the general features of BB structure are conserved across a wide range of species and cell types, there are many examples of dramatic divergence (reviewed in (Gomes Pereira et al., 2021; Jana, 2021; Kumar and Reiter, 2021; LeGuennec et al., 2021). A number of centrioles from various species have been studied using electron tomography under cryogenic conditions (*e.g.*,(Greenan et al., 2018; Guichard et al., 2010; Guichard and Gönczy, 2016; Le Guennec et al., 2020; Li et al., 2012; Li et al., 2019)). A change in diameter from ∼230nm to 220 nm from triplet to incomplete triplet was previously observed in a cryo-ET study of Chinese hamster ovary (CHO) cell centrioles (Greenan et al., 2018), but in that case, the incomplete triplet was reported to be located in the distal centriole instead of in the middle as observed here. In some studies on mammalian centrioles, twist angle variations along the axis of up to 25 degrees have been observed, but in contrast to what we observe in the murine photoreceptor BB, the diameter of the basal body barrel was reported to be constant along its length, unless treated with EDTA (Anderson, 1972; Paintrand et al., 1992), suggesting a role for divalent cations. In basal bodies of motile cilia of *Chlamydomonas reinhardtii*, the centriole barrel was observed to have a constant value of 260 nm throughout its length (Li et al., 2012). However, in a more recent study comparing multiple species centrioles (Le Guennec et al., 2020), the diameter was seen to change from proximal to distal, and in *C. reinhardtii*, the diameter was seen to be greatest in the mid-region of the centrioles. These diameter variations could be due to the triplet-to-doublet transition, changes in twist angles, and/or the change from A-C linker to inner scaffold.

### Non-tubulin complexes: Microtubule Inner Proteins (MIPS) and inter-MT connections

We observe clear densities corresponding to non-tubulin proteins in the lumen of the centriole MT. Although these have been observed in multiple cell types, their positions in motile cilia appear to vary from those in photoreceptors. In most cell types, the molecular identity of these proteins is unknown. The A09/10 MIP has been observed in other centrioles (Greenan et al., 2020), and has been identified in motile cilia axonemes (Ichikawa et al., 2017; Li et al., 2022). This MIP is well conserved throughout species and is seen in our centriole maps as well as the maps we have produced of the A-tubule in the CC. The proteins CFAP53, CFAP161, SPAG8, NME7, CFAP141 and CFAP95, which are found at the A09/A10 seam, are thought to be this MIP (Gui et al., 2021). The C01/C02 bound MIPs were not observed in the triplets with the incomplete C-tubule. The proteins RIB72A and RIB72B have been identified as components of MIPs in the A-tubule in motile cilia of protists (Stoddard et al., 2018). The related mammalian protein EFHC1 (EF-hand-containing protein 1)/myoclonin is a microtubule-associated protein identified as a ciliopathy protein linked to myoclonic epilepsy and present in motile cilia; however, it is not present in mammalian neurons (Suzuki et al., 2020). There is no current information on the proteins making up the MIPs in the photoreceptor BB, though it is likely they are highly conserved because 1) the similar shape and placement of the MIPs observed between previous mammalian maps and the photoreceptor centriole, and 2) the fact that mutations in many of these MIPs cause ciliopathies, which are often associated with blinding disease e.g., CFAP20 (Chrystal et al., 2022).

Also uncertain are the identities of the proteins extending into the lumen of the MT bundle, such as the pinhead and those making up the inter-MT connections, the A-C linker and the inner scaffold. CEP135 has been proposed as a major component of the pinhead, but a definitive determination has not been made (reviewed in (Gonczy and Hatzopoulos, 2019)).The inner scaffold has been described in centrioles from multiple species and has been hypothesized to act as the glue that maintains centriole cohesion and strength under compressive forces (Le Guennec et al., 2020). It has been proposed that the proteins that make up this inner scaffold structure, based on expansion microscopy in human cell lines, include POC1B, FAM161A, POC5, and Centrin-2 (Le Guennec et al., 2020). Since mutations in these proteins are associated with retinal degenerative diseases in humans (https://web.sph.uth.edu/RetNet/disease.htm#10.205d) or animal models (Mercey et al., 2022; Ying et al., 2019), it is possible they form the inner scaffold of centrioles in photoreceptors as well. This structure also resembles the inner scaffold circle observed in conventional TEM throughout the length of the CC. Recently, POC5, centrin, and FAM161A were reported to be localized within the MTs of photoreceptor CC (Mercey et al., 2022), and centrins have long been known to localize within the MTs along the length of photoreceptor CC (Potter et al., 2021; Robichaux et al., 2019; Wolfrum and Salisbury, 1998), consistent with these proteins participating in the inner scaffold. Further work will need to be done to examine where exactly these proteins localize and to determine the structures of the associated protein complexes. An additional non-tubulin complex is the “inner junction” found adjacent to PF10 (Linck et al., 2014) of MT B. This complex has recently been identified as containing the proteins FAP126, FAP106 and FAP276 in motile cilia from protists (Khalifa et al., 2020), and was previously reported to be made up of PACRG and CFAP20 (Dymek et al., 2019). Defining the proteins that directly interact with these at the “stem” of Arm A would be of great interest.

### Limitations of our maps and model

Here we report three distinct compartments in the centrioles of the photoreceptor cilium, with the transition to MTDs occurring within the centriole, providing the template for extension of the doublet-containing axoneme. There is a substantial mid-region with incomplete MTT, but the transition to doublets is clearly complete within the distal centriole, well proximal to the base of the axoneme. It is not clear if this is the general case for mammalian sensory cilia or a special feature of a subset, including those in photoreceptors. We have not included distal or subdistal appendages in our models, which have been built from subtomogram average maps of limited volumes and are based primarily on the MT. These structures, which have been extensively studied in other cell types (Bowler et al., 2019; Chang et al., 2023) are visible in our maps, but would require much larger volumes in the boxing step carried out prior to subtomogram averaging.

### New insights into Y-links and ciliary necklace

The Y-link-containing CC is close to 1 µm in length, considerably longer than typical ciliary transition zones. Although our subtomogram average of this structure is quite noisy, our data provide new insights into the common features of what appears to be a somewhat heterogeneous structure which appears at positions corresponding to the 9-fold positions of the MTD. One of the features that is common among the diverse Y-links within our samples are their connections to the transmembrane portion of the ciliary necklace beads, which appear to be less heterogenous in structure, but somewhat more irregular in their placement on the surface of the ciliary membranes. With sub-tomogram averaging and the increased number of Y-links and ciliary necklace beads in each CC as compared to other transition zones, we were able to, for the first time, resolve these structures to 30-38Å resolution, and to provide strong support to the long-proposed hypothesis that these form one large complex structure extending from the MT bundle through the membrane to its surface. The protein composition of these structures remain uncertain and it is not clear what their functional roles or dynamical properties are. Combining electron microscopic techniques with superresolution fluorescence, proteomics and genetics should clarify these questions in the future.

## Materials and Methods

### Animals

C57BL/6 WT mice, aged 4-6 weeks, were used for this study. All procedures were approved by the Baylor College of Medicine Institutional Animal Care and Use Committee and adhered to the Association of Research in Vision and Ophthalmology (ARVO) guidelines for the humane treatment and ethical use of animals for vision research.

### ROS purification

All mice were maintained in a 12/12 hour light (400 Lux)/dark cycle. Wild-type C57BL/6J were purchased from Jackson lab (Bar Harbor, ME). 1-month C57BL/6J mice used for tissue samples were euthanized by CO_2_ inhalation prior to dissection following American Association for Laboratory Animal Science protocols.

Preparation of purified mouse ROS was modified from (Wensel and Gilliam, 2015). Briefly, to isolate rod outer segments with CC and portions of inner segment attached, retinas were dissected under dim red light and placed in 200 μl Ringer’s buffer (10 mM HEPES, 130 mM NaCl, 3.6 mM KCl, 1.2 mM MgCl_2_, 1.2 mM CaCl_2_, 0.02 mM EDTA, pH 7.4) with 8% (v/v) OptiPrep® (iodixanol, Sigma). Retinas were pipetted up and down with a 200-µl wide orifice tip 50 times, and then centrifuged at 400 × *g* for 2 min at room temperature. The supernatants containing ROS were collected. The process was repeated 4-5 times. All ROS was pooled and loaded onto the top of a gradient of 10, 15, 20, 25 and 30% (v/v) OptiPrep step-gradient and centrifuged for 60 min at 19,210 × *g* at 4 °C in a TLS-55 rotor (Beckman Coulter). The ROS band was collected with an 18G needle, diluted with Ringer’s buffer to 3ml, and pelleted in a TLS-55 rotor for 30 min at 59,825 × *g* at 4 °C. The ROS pellet was resuspended in Ringer’s buffer for cryo-electron microscopy.

### Cryo-ET

Isolated rod outer segments were processed for CryoET as described previously (Gilliam et al., 2012; Robichaux et al., 2019; Wensel and Gilliam, 2015). Briefly, isolated ROS were mixed with BSA-stabilized 15 nm fiducial gold (2:1), and 2.5-3 μL of the mixture was deposited on freshly glow-discharged 3.5/1 200 mesh Quantifoil carbon-coated holey grids. Samples were allowed to settle for 15 s prior to blotting from either the front or back side back-side and plunge-frozen in liquid ethane using a Vitrobot Mark III automated plunge-freezing device. The frozen-hydrated sample was imaged on a Polara G2 electron microscope (FEI Company), equipped with a field emission gun, operated at 300 kV using a direct electron detector camera (Gatan K2 Summit).

Single-tilt image series were automatically collected using SerialEM (Mastronarde, 2005) software at a defocus range of 8-10 μm and a magnification of 9,400 x (equivalent to 4.5 Å/pixel). The total electron dose per tomogram was ∼50-70 e/Å^2^ for 35 tilt images, covering an angular range −51° to +51° with 3° increments (±51°, 3° increment). Each tilt contained. At each tilt angle “movies” of ∼8 frame were collected and Motioncorr (Li et al., 2013) was used to correct the image drift within each tilt before merging them. Alignment and 3D reconstruction were performed automatically using the workflow in EMAN2 software (Chen et al., 2019).

### Subtomogram averaging and correspondence analysis

Structural domains within centrioles and connecting cilia (CC) were visualized and examined longitudinally using IMOD software (Mastronarde, 1997, 2017). Centriole reconstructions were divided into three regions: the proximal region (0-170 nm), containing complete triplets with intact C-tubules, the mid-region (170-340 nm), containing incomplete triplets with partial C-tubules, and the distal region (340-400 nm), containing doublets. The CC (400 nm-CC), containing doublets, was analyzed separately. Sub-volumes from these regions were extracted and assigned to different groups. Subsequently, subtomogram averaging was performed for each group using the *EMAN2* package (Chen et al., 2019).

From 25 centrioles (16 tomograms), ∼1800 sub-volumes from the proximal region, ∼1600 sub-volumes from the middle region, and ∼400 sub-volumes from the distal region, with dimensions of 100 x 100 x 100 nm, were boxed and extracted. The starting models within each group (low-pass filtered to 60 Å) were first generated using *EMAN2* without applying any symmetry, followed by iterative subtomogram refinements performed by *EMAN2* using the corresponding starting models. Resolutions for the refined maps were determined using the gold-standard method of splitting particles into two groups and measuring their Fourier shell correlation (FSC) (FSC=0.143 criterion).For the refinement of triplets, we also used the average triplet structures from *CHO* centrioles (proximal, emd-7776, distal, emd-7777 (Greenan et al., 2018)) as starting models. After a few iterations of refinement, the two different approaches resulted in the consistent/nearly identical maps for both triplets. The final averaged structures have a resolution of 33 Å for both triplets and 40 Å for the doublet. For the doublet averages from CC, ∼1800 sub-volumes with a dimension of 100 x 100 x100 nm was boxed out and yielded a final averaged map at ∼30 Å resolution after several iterative subtomogram refinements by EMAN2.

To further investigate the structure of doublets with “Y-links”, ∼1600 sub-volumes of a microtubule doublet with membrane-directed densities along the CC were boxed and extracted out with a dimension of 100 x 100 x 100 nm. After several iterative subtomogram refinement using the average doublet structure from CC as the initial model, the final model map achieved ∼30 Å resolution. The resulting map had a short density attached to the MTD, but could not be unambiguously interpreted with regard to the orientation (luminal vs. membrane side of MT).

To help identify the luminal vs. membrane side of doublets with Y-links, a single tomogram with longitudinal Y-links spacings of ∼38 nm was selected (Fig. 7A). 150 sub-volumes of microtubule doublets with membrane-directed densities along the CC were boxed out with a dimension of 100 x 100 x 100 nm (including three 32 nm repeats). After several rounds of iterative refinement using EMAN2, a map with ∼38 nm Y-links spacing was obtained, in which the Y-links with 32 nm spacing were clearly visible on the membrane-directed side (Fig. 7A). This map was used for fitting into the rotationally symmetrized CC map as shown in Fig. 7.

For the average of ciliary necklace beads, ∼400 sub-volumes with a dimension of 50 x 50 x 50 nm were boxed along the plasma membrane of ciliary neck and CC regions. After the starting model and iterative refinement were performed by *EMAN*2, the final map achieved a resolution of ∼38 Å.

9-fold averaged maps were used for microtubule triplet/doublet angular twist analysis and “Y-link” visualization in the CC. The centrioles and the cilia with the least distortion of the diameter were picked and visualized *by IMOD* and *Chimera*, and 9-fold symmetry average was applied in *EMAN2* as previously described (Robichaux et al., 2019). To increase the contrast, some of the maps (Fig. 3, Supplementary Fig. S1, Fig. 4G-H) are presented in 4*xbinned* format with a voxel size of 17.8 Å in EMAN2.

### Model building and visualization

The centriole models from the proximal to the distal end as shown in the figures (Figs.2, 4, 7, S1, S6) were built by fitting averaged triplets and doublets manually into a 9-fold symmetry map in UCSF *Chimera* (Pettersen et al., 2004) and then optimizing the fitting using the built-in “*fit in map*” tool. The models for the whole cilia (BB+CC) (Fig. 3) were built by repeatedly fitting the averaged triplets (containing 3-tubulin heterodimers in length) and the doublets back into a raw tomogram. The resulting maps were displayed/presented in 4*xbinned* format in EMAN2. Fitting of averaged structures of MTD with attached Y-links and of ciliary necklace beads was carried out with both raw tomograms and with 9-fold averaged maps. All fitting and superpositions of ciliary necklace beads in TEM images and subtomogram averages, as well as tomographic reconstructions and 3D surface rendering of sub-tomogram averages, were generated and visualized using IMOD (Mastronarde and Held, 2017) and UCSF Chimera (Pettersen et al., 2004) (http://www.rbvi.ucsf.edu/chimera).

### Ambient Temperature Transmission Electron Microscopy

Mice were euthanized under deep anesthesia by transcardial perfusion with fixative (2% PFA, 2% glutaraldehyde, 3.4 mM CaCl_2_ in 0.2 M HEPES, pH 7.4). Eyes were enucleated, the cornea and lens were removed, and the eye cups were placed in fixative (2% PFA, 2% glutaraldehyde, 3.4 mM CaCl2 in 0.2 M HEPES, pH 7.4) for 2 hours, rocking at room temperature. The eyecups were prepared using a similar protocol to that described previously (Potter et al., 2021). Briefly, the eyecups were embedded in 4% agarose from which 150 μm vibratome sections were cut. These sections were subsequently stained (rocking at room temperature) with 1% Tannic Acid with 0.5% Saponin in 0.1 M HEPES, pH 7.4, for 1 hour followed by 1% uranyl acetate in 0.2 M Maleate Buffer, pH 6.0, for 1 hour. The sections were dehydrated in a series of 15 minute ethanol washes (50%, 70%, 90%, 100%, 100%), followed by infiltration with Ultra Bed Epoxy Resin (Electron Microscopy Sciences). The sections were embedded in resin between 2 ACLAR sheets sandwiched between glass slides in a 60°C oven for 48 hours. At 24 hours, the top slide and ACLAR sheet were removed, and resin blocks in BEEM capsules (Electron Microscopy Sciences) were stamped onto each section to allow for polymerization the following 24 hours.

For increased resolution of the Y-links, mice were euthanized by CO_2_ asphyxiation followed by cervical dislocation and after enucleation, retina were dissociated from eyecups and incubated, rolling, in 0.1% TritonX-100 in 1xPBS for 1 hour at 4°C. After rinsing with 1xPBS, they were fixed for 2 hours in the same fixative as above. The retinas were directly stained, dehydrated, and infiltrated as above in half-dram (1.35 mL) glass vials. They were embedded in BEEM flat embedding molds in a 60°C oven for 48 hours.

Ultrathin sections (50-70nm) were cut on a Leica UC7 ultramicratome and were placed on copper grids and poststained in 1.2% uranyl acetate in MilliQ water for 6 minutes, followed by staining in Sato’s lead (a solution of 1% lead acetate, 1% lead nitrate, and 1% lead citrate; all from Electron Microscopy Sciences) for 2 minutes. Grids were imaged on either a JEOL 1400 Plus electron microscope with an AMT XR-16 mid-mount 16-megapixel digital camera or on a JEOL JEM-1400Flash 120 kV TEM with a high-contrast pole piece and a 15 megapixel AMT NanoSprint15 sCMOS camera. For each microscope, AMT software was used for image acquisition and images were subsequently cropped with slight contrast adjustments in FIJI//ImageJ (Schneider et al., 2012).

### Twist Angle Measurements

For the estimation of twist angle, two methods were used. In the first method (β-angle), nonagon vertices were manually placed in the center of each B-tubule and a midline drawn through the centers of the tubules of each triplet/doublet, and then the angle between each triplet/doublet (by midline) and its closest nonagon edge was measured. This method was used for the angular estimation in centriole maps (BB) with and without imposition of 9-fold symmetry, and also for the whole cilium raw map (BB+CC). In the second method (α-angle), nonagon vertices were placed in the center of A-tubules rather than B-tubules. This method was used only for the angular estimation along the whole cilia length (BB+CC) for comparison with motile cilia (Greenan et al. JCB, 2019). The angles were measured and averaged from the cross-section every 90 nm and displayed as a function of distance throughout the cilium axis (proximal-distal end).

## Data Availability

Cryo-ET data and calculated maps have been deposited with the Electron Microscopy Data Base under EMDB entry IDs EMD-41812, 41813, and 41814. All other data are available on request.

## Acknowledgements

Funding was provided by grants from the NIH (R01-EY026545, R01-EY031949, F32-EY-031574, R21MH125285, and the Welch Foundation (Q0035). Thanks to the BCM cryo-EM core, the cryo-EM core at the University of Texas Health Sciences Center, Houston, Lita Duraine of the TEM core at the Neurological Research Institute, Houston. The authors declare no competing financial interests.

## Author Contributions

ZZ designed and executed all cryoET imaging and subsequent image processing and analysis. FH performed ROS isolation. ARM designed and executed all procedures and analysis related to ambient temperature TEM. MC provided advice and performed some analyses related to sub-tomogram averaging. ZZ and ARM designed and prepared all figures with edits by other authors. ARM prepared the initial manuscript draft, which was edited by ARM, ZZ, MAA and TGW. TGW supervised research and secured funding.

## Abbreviations

BB: basal body
CC: connecting cilium
Cryo-ET: cryo-electron tomography
DMT: doublet microtubule
OS: outer segment
ROS: rod outer segments
TEM: transmission electron microscopy
TMT: triplet microtubule
TZ: transition zone

## Supplementary figures

**Figure S1.**
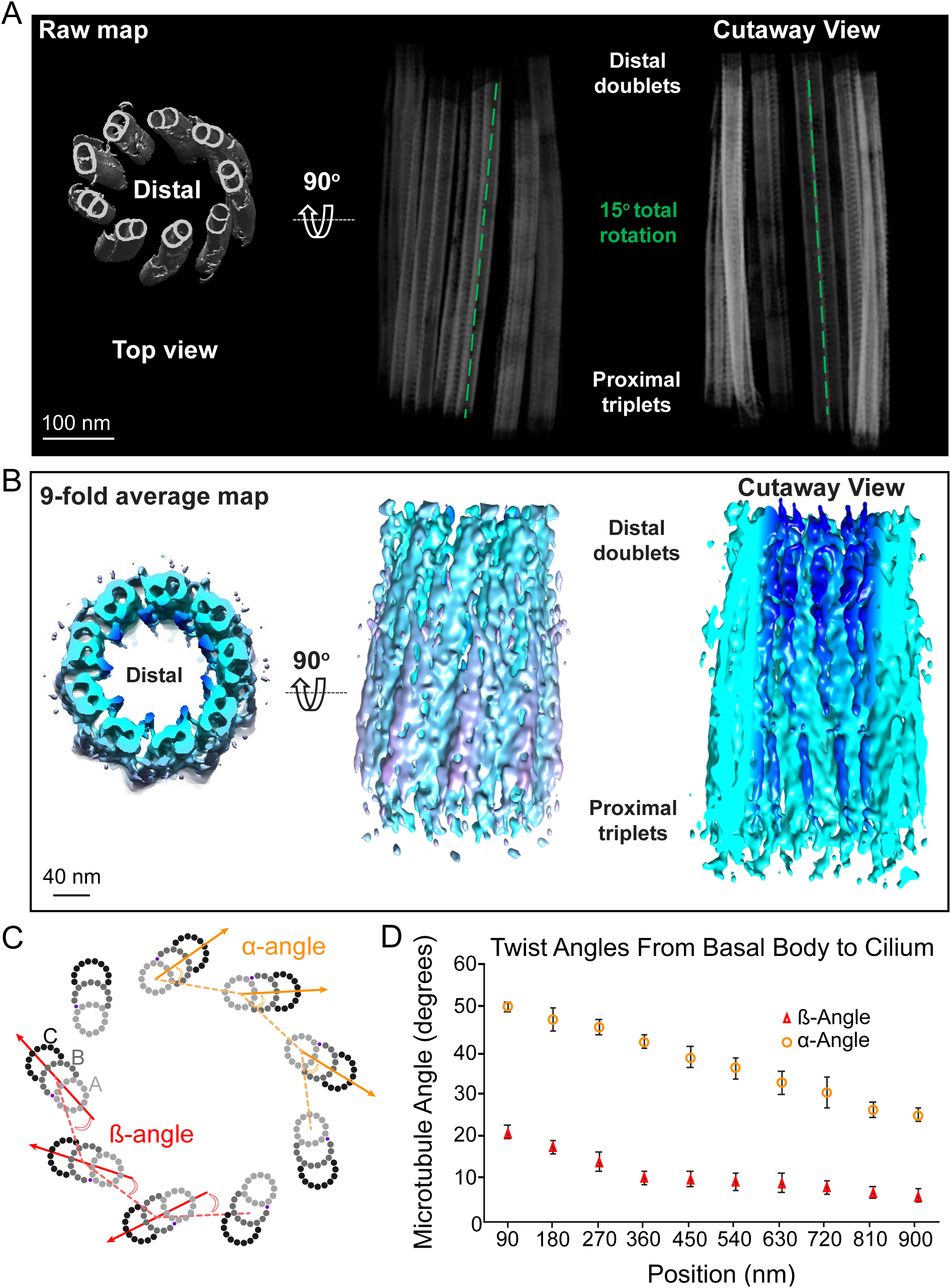
Axial and cross-sectional twist angles of centriole and CC. (A) Projection views of the raw tomogram reconstruction of the centriole from the top (showing the distal end of the centriole) and side views. To the far right is a cutaway view. The side views illustrate the gradual twist of the entire centriole (highlighted with cyan (left) and green (right) dashed lines). (B) The 9-fold symmetrized map of the centriole displayed as for the raw map in (A), except for radial distance-coded color scheme and surface view. The diameters are indicated in (B). (C) Schematic diagram of centriole MTTs and how the ß-angles and α-angles are measured (see main Figure 3). Note that for 9-fold symmetry, these two angles are related as:

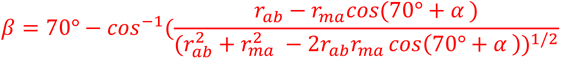

Where *r*_*ab*_ is the distance between the centers of the A and B MT, *r*_*ma*_ is the distance from the center (middle) of the MT bundle to the center of the A MT. (D) Variation of the triplet and doublet twist angles along the entire cilium. Points and error bars represent means ± S.D.

**Figure S2.**
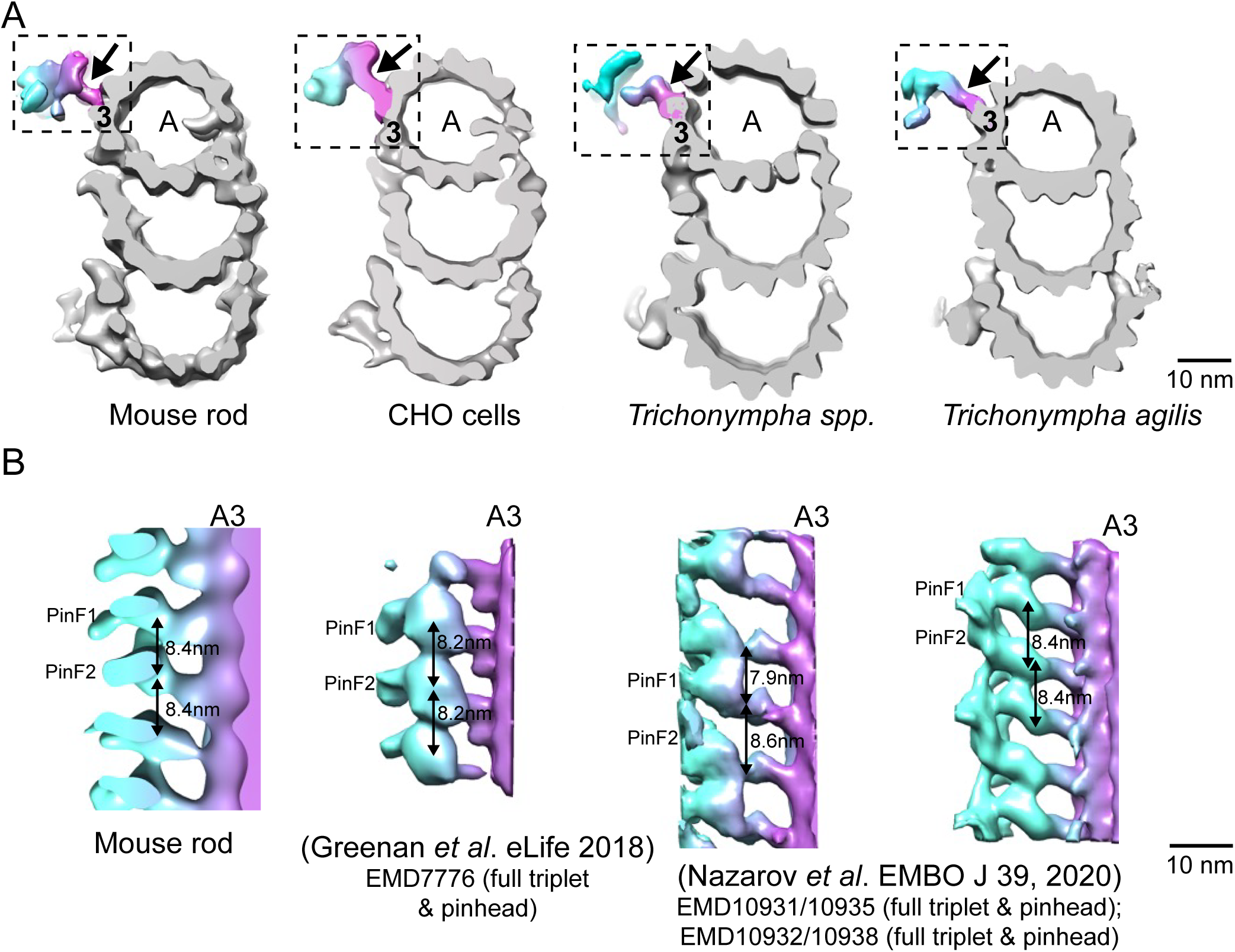
Comparison of pinhead architectures of complete triplets from different species. (A) Cross-sectional views of MTTs from mouse photoreceptor, Chinese hamster ovary (CHO) cells (Li et al., 2019), *Trichonympha Spp.* (generic term for a groups of termite gut protists) *and Trichonympha agilis (Nazarov et al., 2020).* All the pinheads (dashed box) connect to A3 protofilament. (B) Longitudinal views of the pinheads (arrows in (A)). The pinheads have similar orientations, with the two pin feet moieties PinF1 and PinF2 extending from A3 PF. In *T. Spp*, the average spacing between PinF1 and PinF2 of adjacent pinheads (7.9 nm) differs from that between PinF1 and PinF2 of the same pinhead (8.6 nm), whereas for the other species these two spacings are the same: 8.4 nm for WT mouse, 8.2 nm for CHO, and 8.4 nm for *T.agilis*.

**Figure S3.**
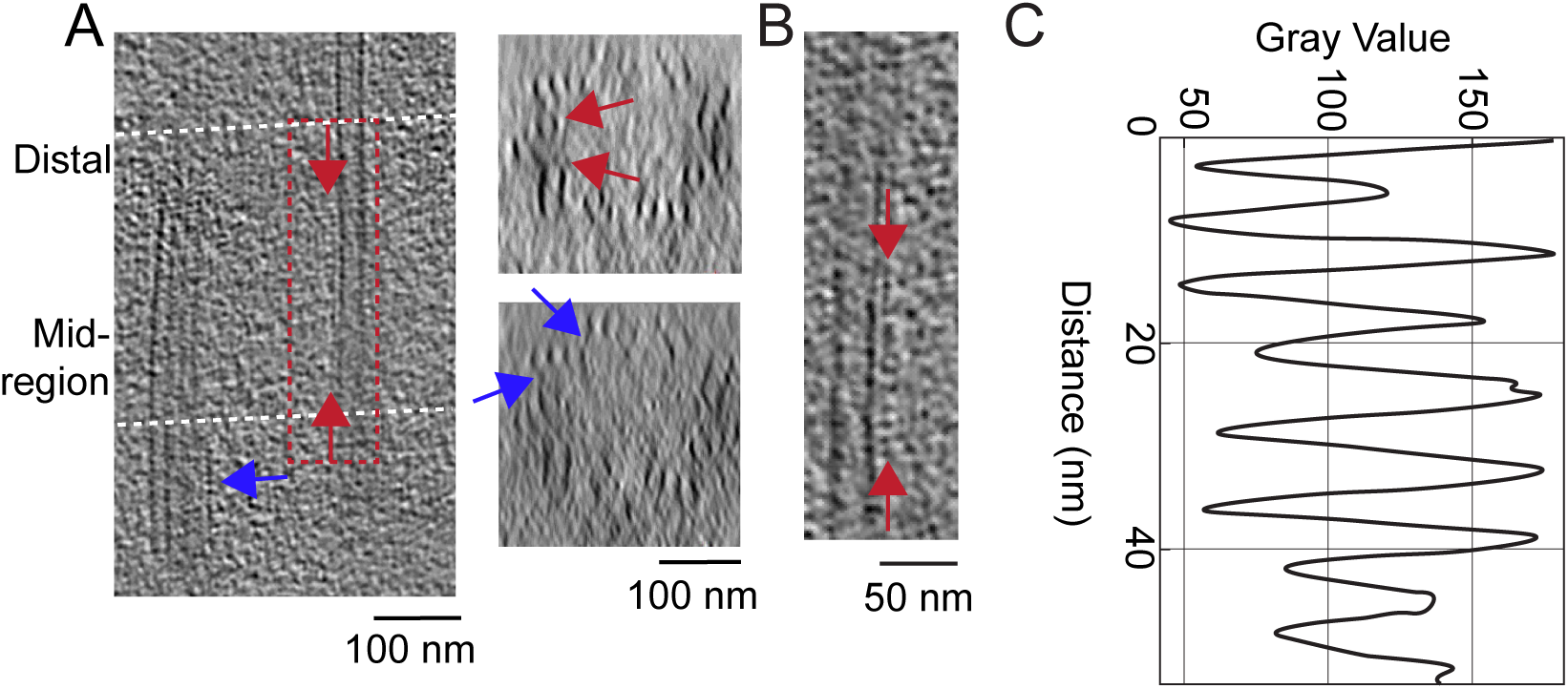
Visualization of the connections between MTTs (by AC-linker) and MTDs (by inner scaffold) in raw tomograms. (A) Section through raw tomogram generated by Cryo-ET of centriole showing the entire structure in longitudinal view. Dashed lines indicate the positions of the cross-sections shown to the right. Arrows indicate the visible inner scaffolding in MTDs (red arrows), and A-C linker connecting MTT in the proximal region (blue arrows).

**Figure S4.**
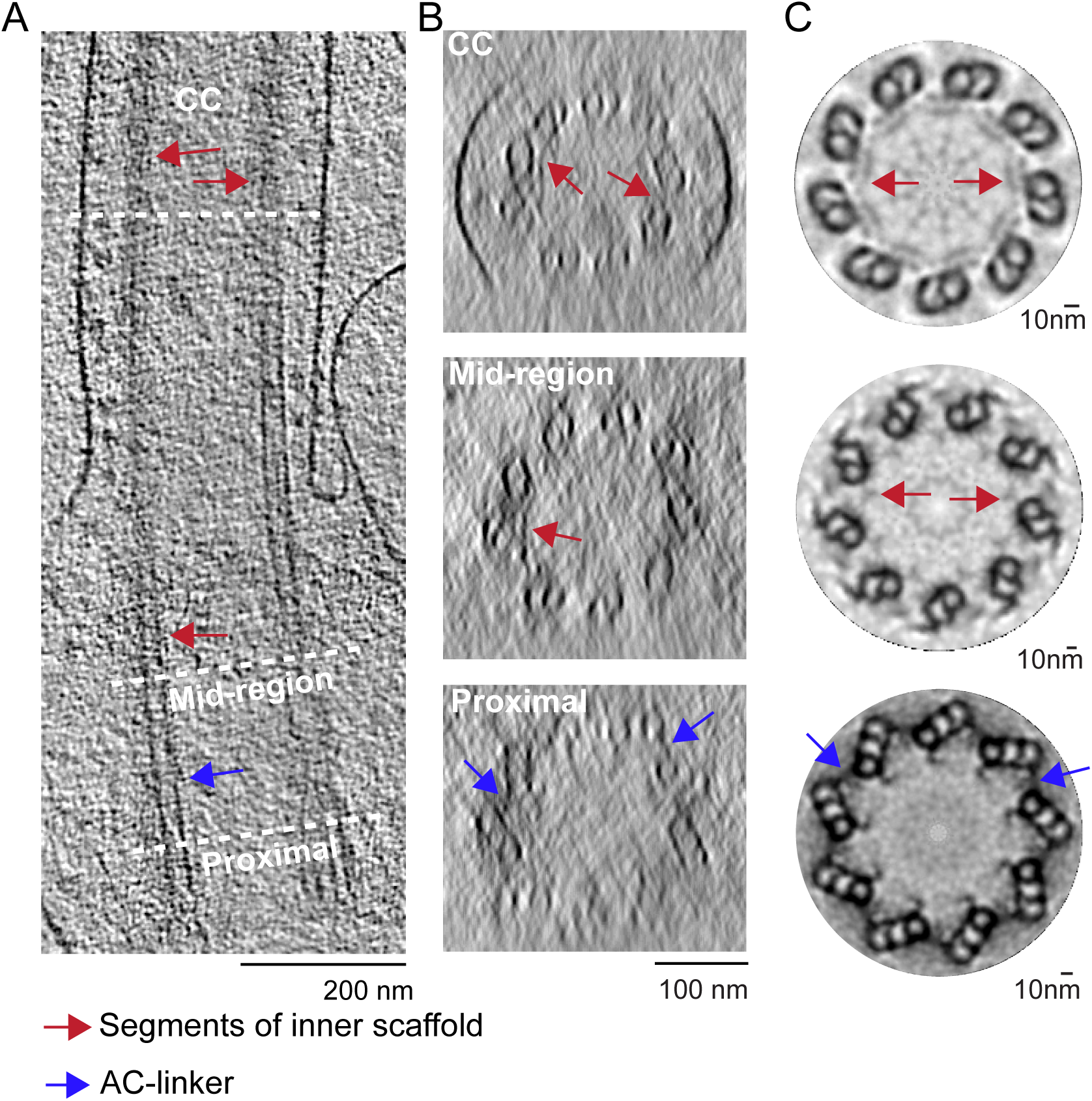
Visualization of the inner scaffold through centriole and CC. (A) Longitudinal section of a raw cryo-tomogram. Dashed lines indicate the positions of the cross sections in (B). (B, C) Cross-sections from the three regions along centriole and CC length, (B) before and (C) after ninefold symmetry averaging. Arrows show the inner scaffold partially attached to the doublets (red arrows) and the A-C linker within adjacent triplets (blue arrows).

**Figure S5.**
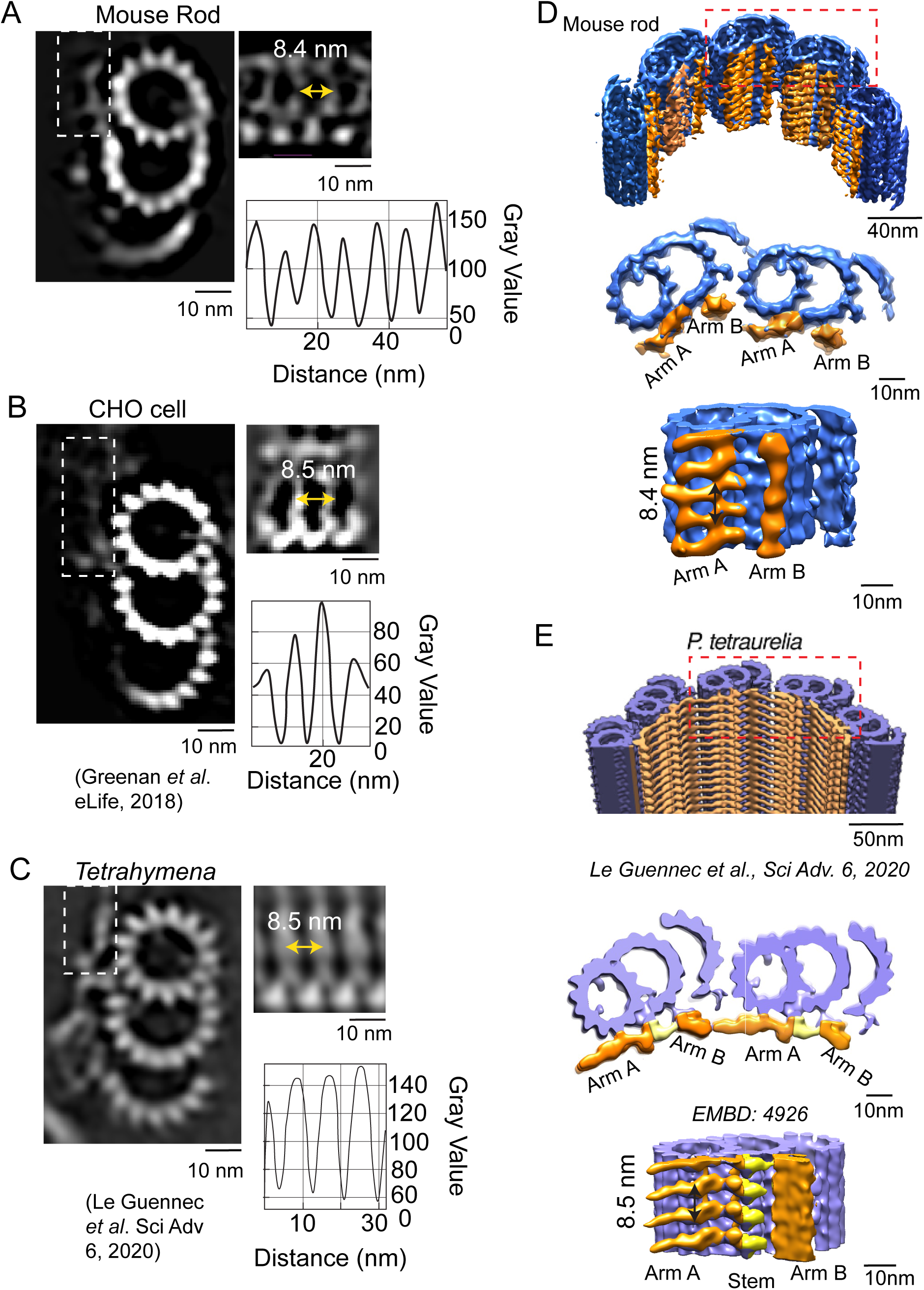
Comparison of inner scaffolds and Arm A-Arm B scaffold in centrioles from different species. (A-C) (Left) Cross-sectional slices from raw tomograms through region at the beginning of the transition from complete to incomplete triplets in (A) WT mouse photoreceptor (this work), (B) CHO cell (Greenan et al., 2018), and (C) *Tetrahymena* (Le Guennec et al., 2020). The regions outlined with dashed-line boxes are shown to the *right* rotated 90° to the right into the plane of the page, and show Arm A extending from A/B junction. The corresponding intensity profiles (*far right*) display similar periodicity among the species (∼8.4 nm and 8.5nm). **(D and E)** Different views (tilted cut away views from the lumen of the MT bundle, and cross-sectional views of the models from mouse rod photoreceptors and *P. tetraurelia* (E) are shown, with incomplete MTT in blue (mouse rod, D) or violet (*P. tetraurelia*, E) and Arm A Arm B shown in gold. A stem density connecting the two is colored yellow in the *P. teraurealia* map, but this feature is much weaker and not visualized at this contour level in the mouse rod (D). The inside views from the centriole interior show the attached inner scaffold with spacing at ∼8.4 nm for mouse photoreceptor (D) and ∼8.5 nm for *P. teraurealia* (E).

